# Transcriptomic similarity guides source selection beyond tissue lineage

**DOI:** 10.64898/2026.10.26.761823

**Authors:** Li Jiang, Jiahong Pang, Bote Luo, Xi’ai Wu, Xin Li, Haojun Zhang

## Abstract

Perturbational and pharmacogenomic maps remain incomplete across cellular contexts, leaving a practical source-selection question before response prediction: which measured context should inform an unmeasured target? We tested whether baseline transcriptomic similarity ranks candidate sources for held-out targets, separating source ranking from aggregation and transfer-uncertainty decisions. Similarity tracked matched-response concordance in LINCS and GSE306429; selected-source utility exceeded random expectation in GSE306429 (ΔU = 0.034; P = 0.002) and in exact-time-matched MIX-Seq tasks (ΔU = 0.046). In the full GDSC2 analysis, the drug-balanced ranking effect was 0.359 (95% CI, 0.337–0.383). In a lineage-restricted analysis of the same eligible subset, within-tissue ranking remained positive at 0.130 (95% CI, 0.119–0.141), retaining 35.8% of the same-subset global effect (0.363; 95% CI, 0.342–0.386). Global transcriptomic top-1 selection exceeded tissue-matched random selection by 0.154 (95% CI, 0.143–0.164). Thus, baseline similarity adds source-ranking information after tissue matching in the evaluated setting, while ranking alone does not resolve whether to aggregate sources or how to quantify transfer uncertainty.

## Introduction

Large perturbational resources connect compounds, genetic perturbations and cell states to transcriptional responses across many cellular backgrounds. The Connectivity Map, L1000 and LINCS expanded this measurement space (Lamb et al., 2006; Subramanian et al., 2017; Keenan et al., 2018). Yet no resource measures every perturbation in every context. When a target response is missing, a measured context may need to serve as its source.

Baseline expression, tissue lineage and genomic state carry information about pharmacogenomic response (Barretina et al., 2012; Ben-David et al., 2018; Chawla et al., 2022; Garnett et al., 2012; Iorio et al., 2016). Response prediction has used expression-based models, cell-line selectivity, transfer learning and drug-response benchmarks in defined settings (Bernett et al., 2026; Carli et al., 2025; Chen et al., 2022; Costello et al., 2014; Feizi et al., 2022; Geeleher et al., 2014; Shimada et al., 2016; Zhang et al., 2015). Large pharmacogenomic comparisons document study-level variability (Haibe-Kains et al., 2013; Safikhani et al., 2017), while molecular alignment can connect tumour and cell-line profiles (Warren et al., 2021). These approaches establish molecular state as a useful predictor or comparator, but they do not directly test which measured source is most useful for a held-out target.

Transcriptomic signature matching has been used to rank compounds for repurposing, and active-learning systems have used transcriptomics to prioritize interventions for testing (DeMeo et al., 2025; Pham et al., 2022). Perturbation studies show shared and context-specific response components (Dixit et al., 2016; McFarland et al., 2020; Replogle et al., 2022; Srivatsan et al., 2020), with gene-regulatory network structure informing the distribution of perturbation effects (Aguirre et al., 2025). Response predictors range from single-cell models to multi-perturbation and foundation-model evaluations (Ahlmann-Eltze et al., 2025; Crawford et al., 2025; Csendes et al., 2025; Kernfeld et al., 2025; Lotfollahi et al., 2019, 2023; Qi et al., 2024; Viñas Torné et al., 2026). These approaches ask which perturbation to test or how to predict a response; source selection asks which already measured context is most useful for a specified target. This comparison requires held-out targets and a candidate pool, with utility defined by the response endpoint.

Here we test whether baseline transcriptomic similarity can rank measured sources for a held-out target, including when candidates are restricted to the target tissue. LINCS and GSE306429 evaluate matched-response association, with GSE306429 also supporting a task-level source-selection test; MIX-Seq provides an exact-time-matched external task evaluation; and GDSC2 extends the decision to pharmacogenomic sensitivity. In GDSC2, the within-tissue ranking estimate is compared with a global comparator on the same eligible subset, and global top-1 selection is compared with tissue-matched random selection. This estimates the incremental ranking information after tissue matching in the evaluated resource, without implying independence from lineage. Source ranking, aggregation and transfer uncertainty are evaluated as separate decisions.

## Results

### Study architecture and estimands

We defined a target as a cellular context with an unmeasured perturbational or pharmacogenomic response and a source as a measured context eligible to inform that target (Fig. 1). Baseline transcriptomic similarity ranked candidate sources. Downstream analyses kept three decisions separate: selecting one source, aggregating several sources and estimating whether transfer should be trusted. This design prevents a positive association or ranking result from being interpreted as evidence for an ensemble rule or an applicability-domain detector.

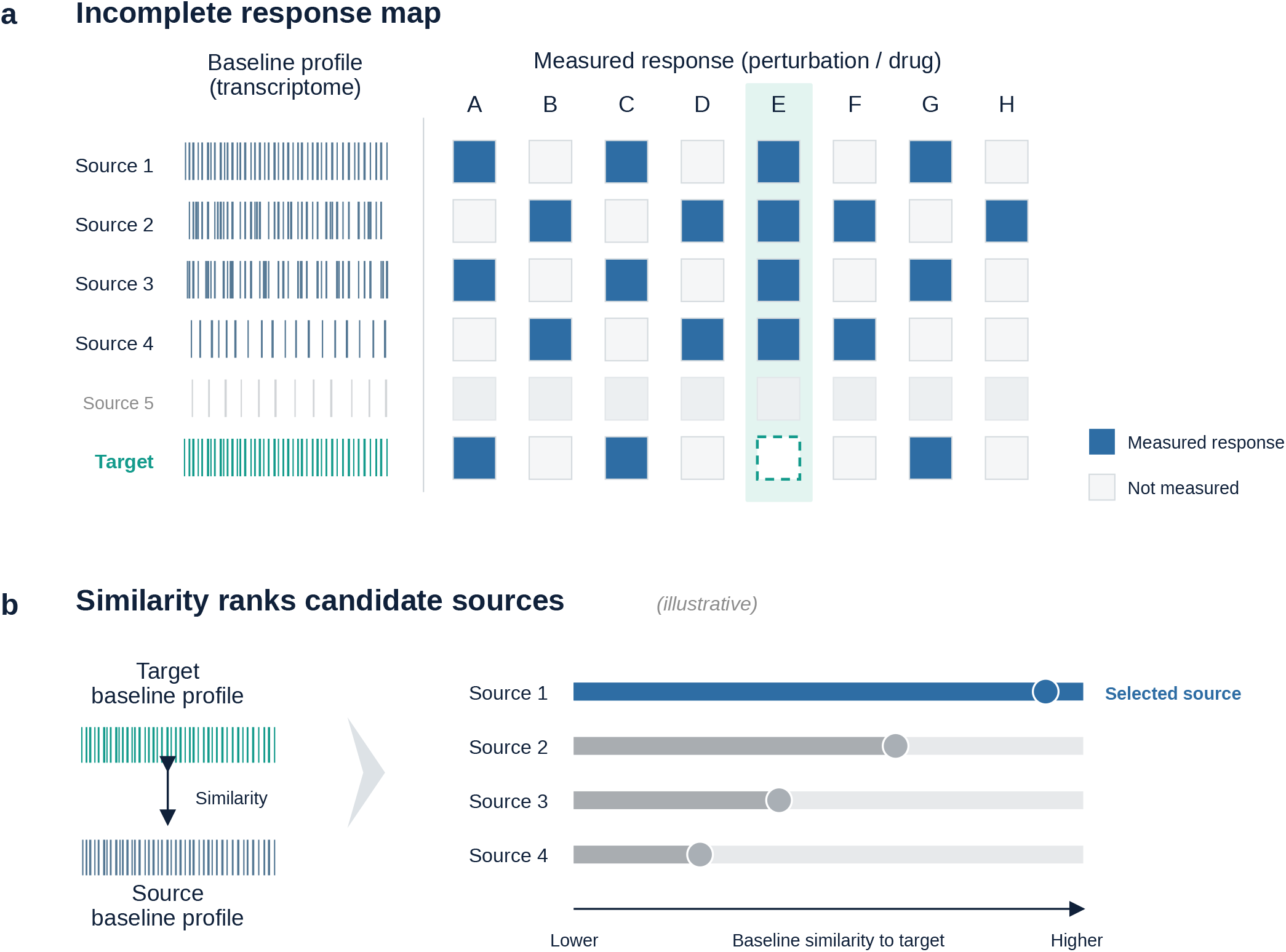

The evidence set included 471,306 LINCS pair-condition records nested within 435 cell pairs, a GSE306429 association analysis and task-level utility analysis, 808 MIX-Seq tasks across cancer-derived backgrounds, and two GDSC2 scopes. The full GDSC2 analysis contained 97,077 target-drug tasks across 295 drugs and 396 exact joint models. The lineage-restricted analysis used a separate eligible subset of 92,324 tasks across 295 drugs, 377 target models and 18 represented tissues. Record counts, pair counts, task counts, drug counts and model counts were retained as distinct units throughout.

### Baseline state carries a reproducible source-ranking signal

In the record-level primary model, response cosine similarity was positively associated with baseline Pearson similarity after adjustment for response-signature strength covariates (beta = 0.255; standard error = 0.006; R2 = 0.327; n = 471,306; empirical P < 0.001). Supporting response outcomes were directionally consistent, including response Pearson similarity, signed direction agreement and signed top-100 overlap.

Collapsing to 435 cell pairs gave a dyadic coefficient of 0.290. Two-way clustering on the two dyad endpoints produced a 95% CI of 0.218-0.363 across 435 dyads and 30 cell lines (Cameron et al., 2011). A separate pair-level node-label permutation test reached its plus-one resolution floor (empirical P < 0.001), consistent with a positive association under dyadic inference.

Baseline similarity was high (median Pearson similarity 0.801), whereas response cosine was more modest (median 0.108; mean 0.131), indicating enrichment for greater response concordance rather than a deterministic match. Leave-one-cell-line-out and leave-one-cell-pair-out estimates remained positive across evaluable groups.

### Directional extension across an independent perturbation resource

We next examined GSE306429, an independent single-cell perturbation resource with matched-DMSO response signatures (Fig. 2). The analysis included 10 eligible backgrounds and 45 eligible pairs. The response catalogue contained 760 eligible background-by-compound-by-dose conditions across 103 compounds and 11 dose values. Pair comparison retained 2,763 pair-condition records, with shared conditions per pair ranging from 4 to 88.

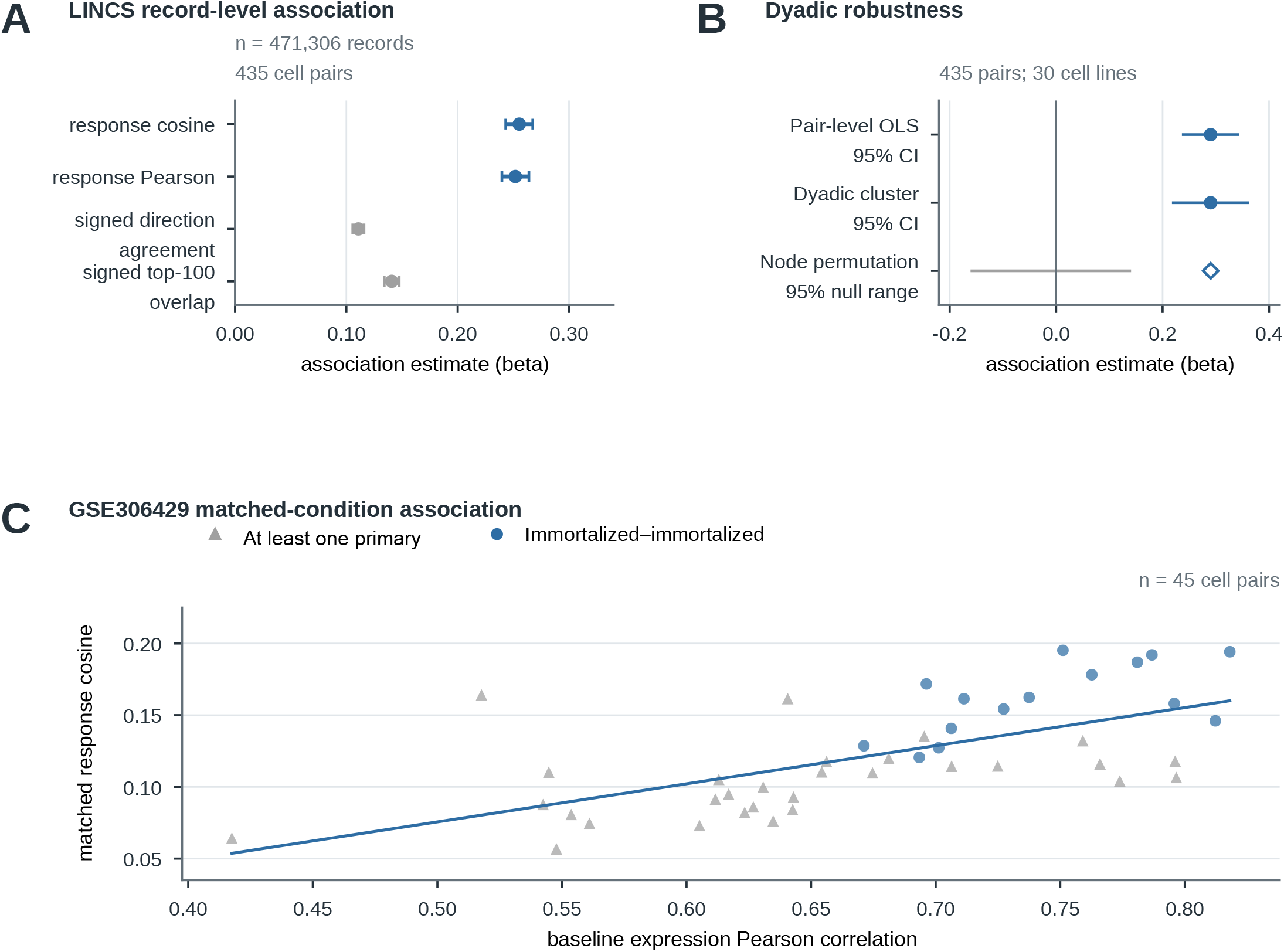

The pair-level estimand was the mean response cosine across exactly matched compound-dose conditions within each pair. The descriptive pair-level coefficient was 0.266, with standardized beta = 0.641 and R2 = 0.411. Shared-node robust inference gave a 95% CI of 0.165-0.366 and P < 0.001. A synchronous node-label permutation test gave empirical P < 0.001, and all 10 leave-one-node-out estimates remained positive. The direction therefore extended beyond the LINCS platform under the matched-condition estimand.

The pair-level analysis asks whether baseline and response similarity covary across background pairs; the task-level analysis asks whether selecting a source improves utility for a held-out target. These analyses use different units.

### Similarity improves individual-source selection in perturbational settings

For each task-level analysis, candidate sources were ranked by baseline similarity, and the highest-similarity source was compared with the mean utility expected from the eligible single-source candidate pool (Fig. 3). Across 736 GSE306429 tasks, the mean selected-minus-random utility difference was 0.034 and the permutation P value was 0.002. The result remained positive under perturbation-balanced bootstrap resampling, with a 95% CI of 0.027-0.042.

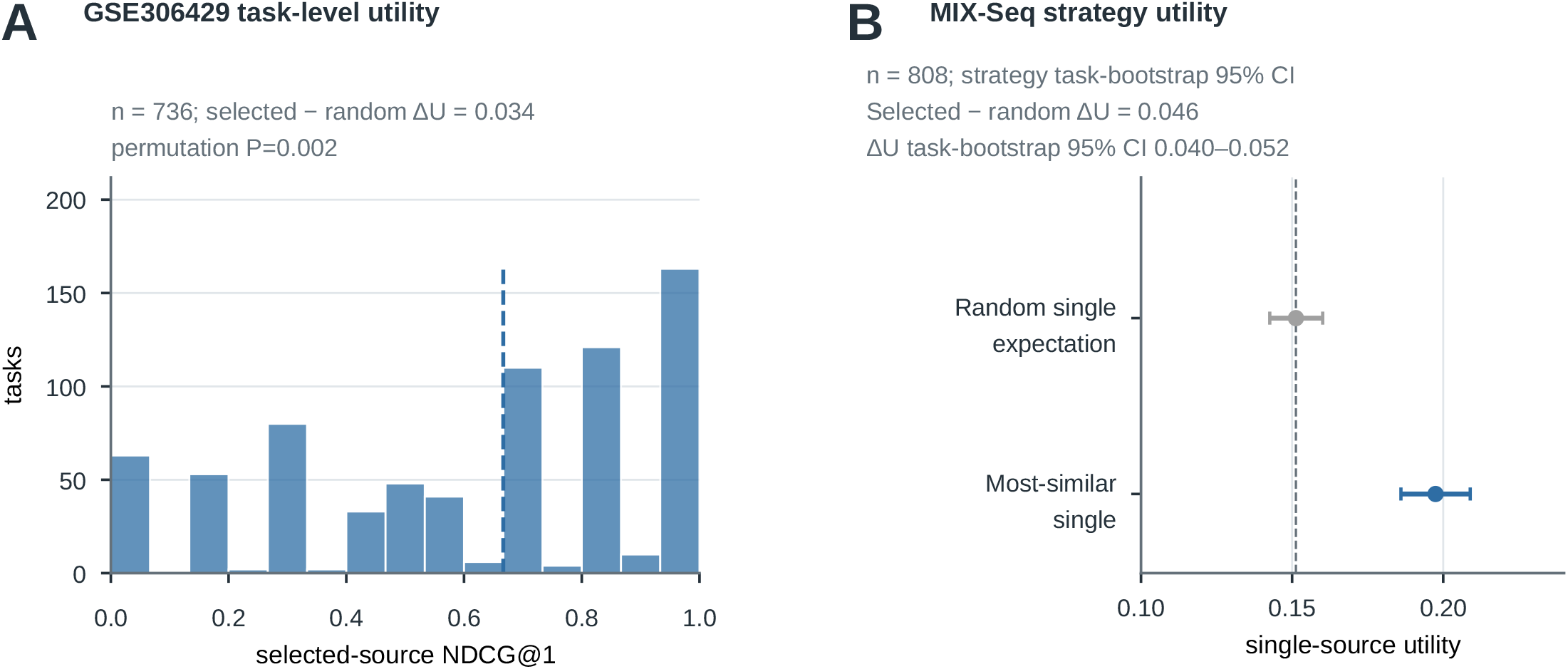

MIX-Seq provided an external cancer-derived evaluation under exact-time baseline matching. Across 808 tasks, the most-similar single-source utility difference was 0.046 (perturbation-balanced 95% CI 0.037-0.056). Availability was concentrated in the evaluated time and compound strata: 4,166 of 54,640 audited baseline requests were unavailable and were excluded without imputation or cross-time substitution. The weighted-minus-unweighted ensemble contrast included zero under perturbation-balanced resampling. These results support bounded individual-source ranking in the evaluated candidate pools, not unrestricted cross-class transfer.

### Full GDSC2 analysis extends ranking to pharmacogenomic sensitivity

The full GDSC2 analysis used official LN IC50 response records and exact SIDM identity matching between response models and RNA-seq profiles (Fig. 4). The analysis covered 97,077 target-drug tasks across 295 eligible drugs and 396 exact joint models. The primary estimate was the equal-weight mean of the 295 drug-level mean single-source utility effects: 0.359 (95% drug-bootstrap CI 0.337-0.383), and all 295 drugs had a positive drug-level mean. The task-weighted mean was 0.363 and is descriptive only; task-level observations were not treated as independent inferential units. A positive utility difference was observed in 69.5% of tasks. Figure 4 therefore addresses pharmacogenomic source ranking only.

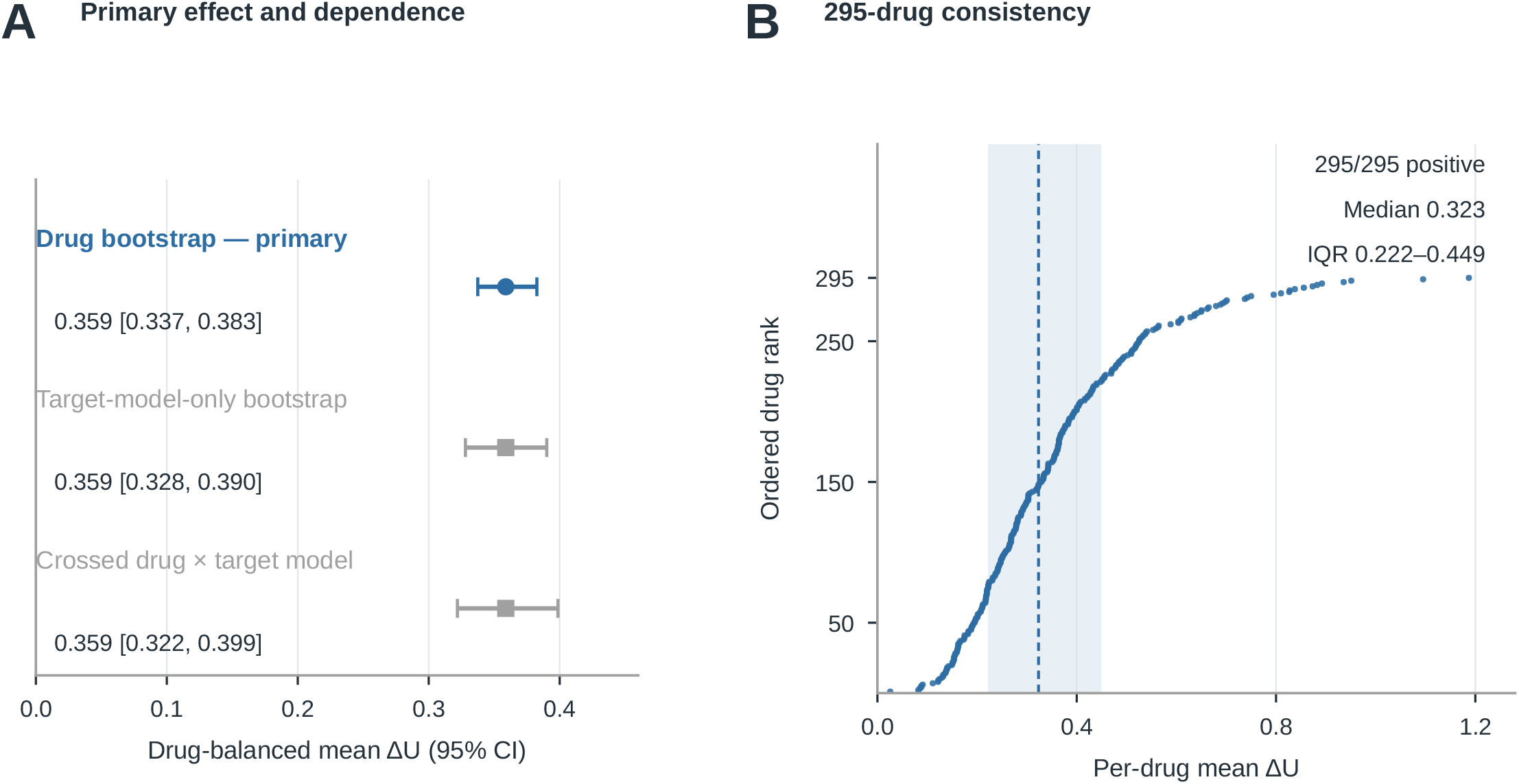

### Ranking persists when candidate sources are restricted to tissue lineage

We next evaluated the lineage-restricted analysis on the same eligible subset of 92,324 target-drug tasks, 295 drugs, 377 target models and 18 tissues (Fig. 5). The global comparator within this subset gave an effect of 0.363 (drug-bootstrap 95% CI 0.342-0.386). Restricting candidate sources to the target tissue gave a positive within-tissue effect of 0.130 (drug-bootstrap 95% CI 0.119-0.141; crossed drug-by-target-model 95% CI 0.101-0.158). The within-tissue effect was 0.358 of the same-subset global effect (95% CI 0.338-0.378), corresponding to 35.8% retention.

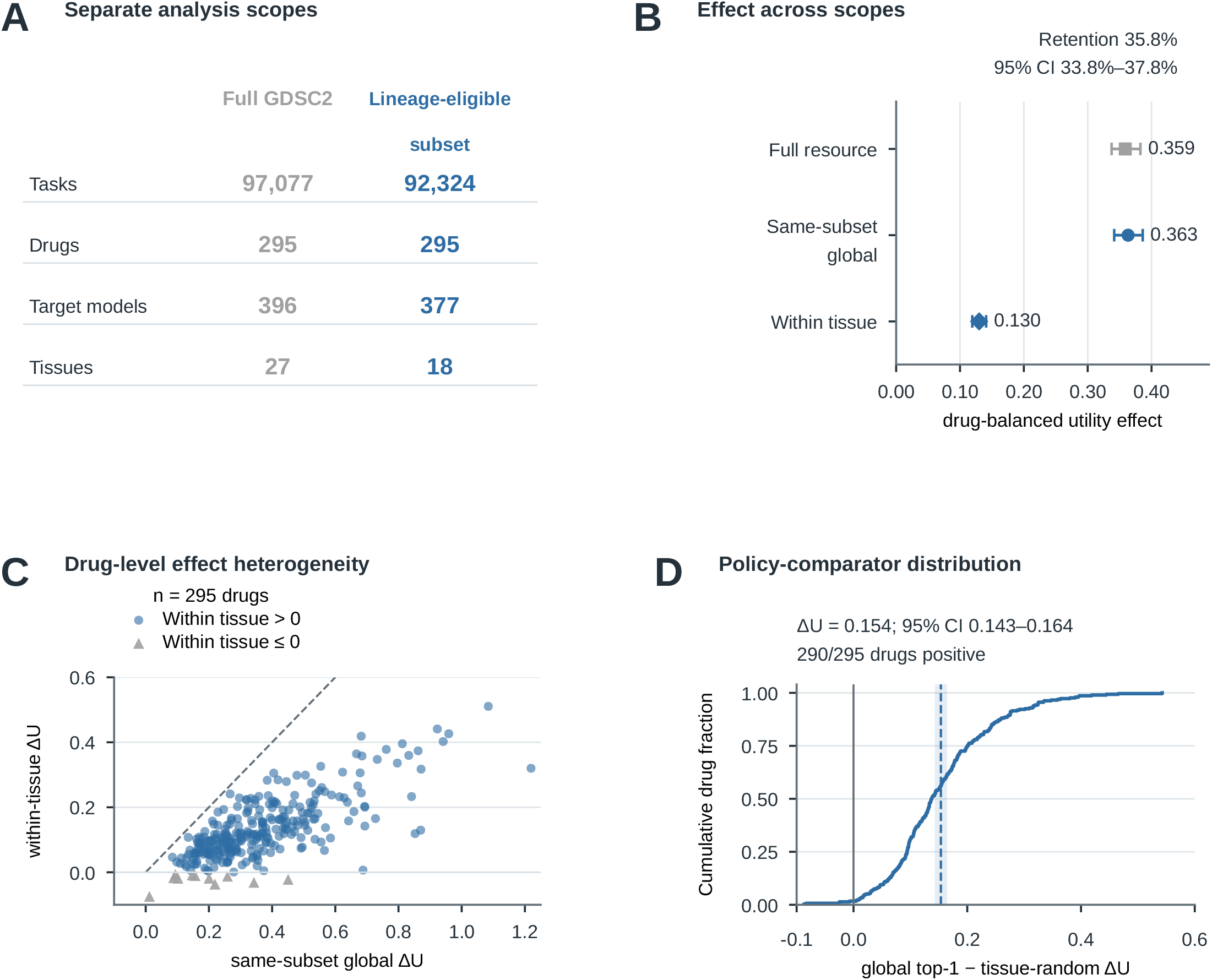

The policy comparison gave the same direction. Global transcriptomic-similarity top-1 selection exceeded a tissue-matched random-source policy by 0.154 (drug-bootstrap 95% CI 0.143-0.164; crossed 95% CI 0.126-0.183). This is a comparison of source-selection policies within the evaluated candidate pools; it is not a causal decomposition of tissue and molecular state. Cancer-type sensitivity analysis gave an effect of 0.091. In a candidate-level adjusted model, the similarity coefficient was 2.054 (crossed 95% CI 1.838-2.282) and the same-tissue coefficient was 0.225 (crossed 95% CI 0.183-0.269). Coverage was 95.1% of reconstructed tasks, 100% of drugs, 95.2% of target models and 66.7% of tissues.

### Ranking, aggregation and transfer uncertainty require separate evidence

We evaluated whether similarity-related quantities could identify tasks in which a selected source would underperform an all-source ensemble (Fig. 6). In the perturbational utility benchmark, equal-weight combination was stronger on average than the selected individual source: the selected-minus-ensemble difference was -0.053 (95% CI -0.071 to -0.034). The same aggregation boundary appeared in GDSC2: the equal-weight mean of 295 per-drug selected-minus-ensemble values was -0.083 (95% drug-bootstrap CI - 0.097 to -0.069), with selected-source utility higher for 66 drugs and ensemble utility higher for 229. The task-weighted GDSC2 mean was -0.081 and is descriptive only, with no corresponding task-level interval.

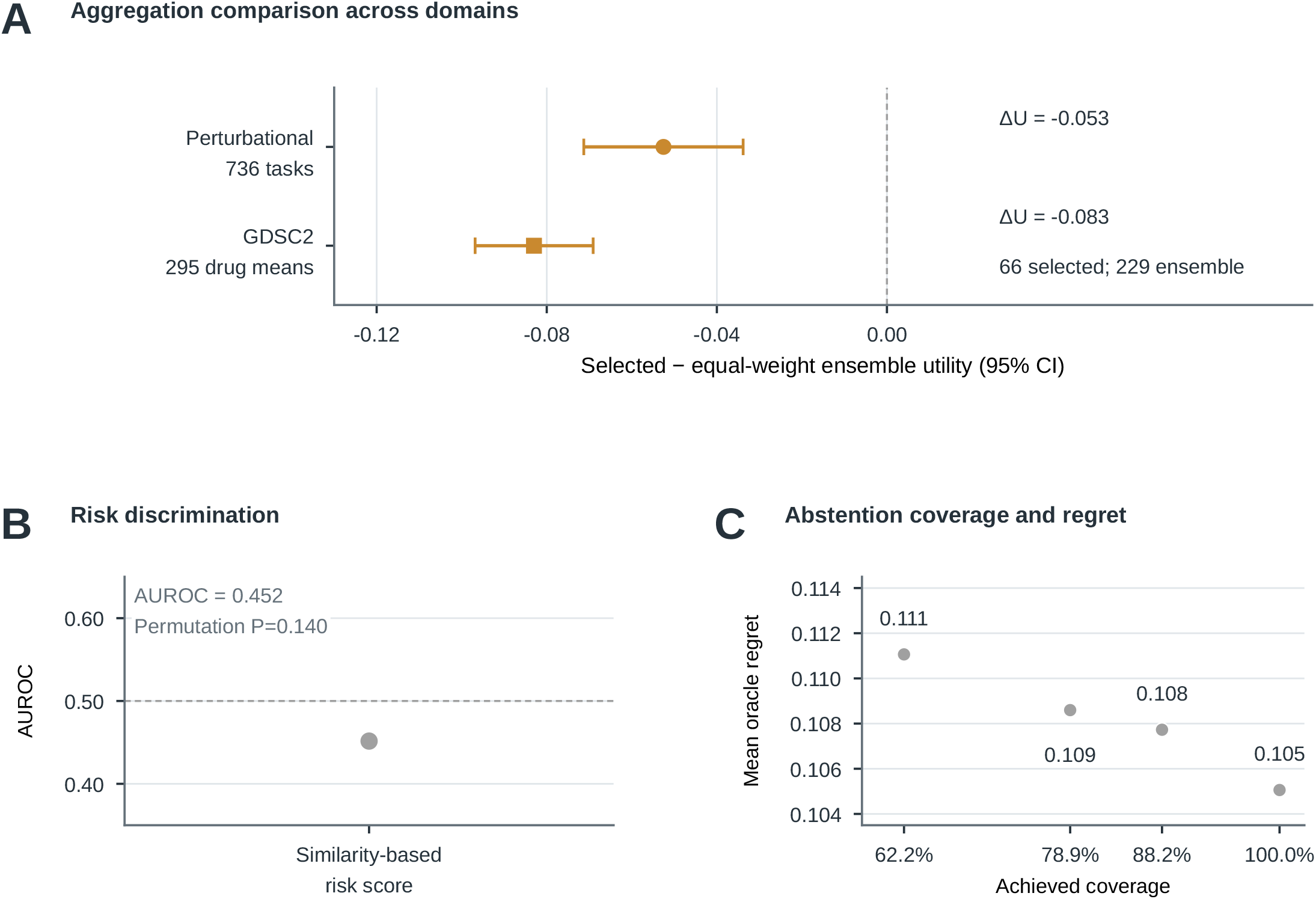

Similarity-based risk discrimination was not supported (AUROC = 0.452; permutation P = 0.140). The abstention analysis is shown at its four frozen coverage points only: regret was 0.105 at full coverage and 0.109 at 78.9% achieved coverage (the prespecified 80% operating point). These endpoints distinguish source ranking from aggregation and transfer uncertainty.

## Discussion

This study identifies source selection as an operational systems-biology decision that precedes response prediction. Across perturbational resources and full GDSC2 pharmacogenomic data, baseline transcriptomic state helped identify which measured context was more informative for an unmeasured target. The lineage-restricted result shows that this information persisted after candidate sources were limited to the target tissue: the within-tissue effect retained 35.8% of the same-subset global effect, and global top-1 selection exceeded a tissue-matched random policy.

The result changes how incomplete molecular maps can be used. Tissue lineage remains a useful organizing variable, but it need not be the only source-selection criterion among measured contexts. A molecular similarity score can serve as a ranking prior that narrows the source pool before a response model is applied. This is a decision function rather than a replacement for direct response prediction, and the within-tissue comparison does not establish molecular independence from lineage.

The ranking signal does not identify when a source ensemble is preferable or when transfer should be withheld. In the perturbational analyses, equal-weight aggregation performed better on average, while the evaluated risk and abstention measures did not support a transfer rule. Combining sources and calibrating uncertainty therefore require their own validation outcomes.

The analysis covers the evaluated cancer-derived resources, response definitions, matching rules, tissues and candidate pools; performance in other biological settings or clinical use remains to be tested.

## Methods

### Study design and analysis units

All task-level source-selection analyses used the same decision structure: for a held-out target, rank eligible sources by baseline similarity and compare the selected source with the mean utility across the candidate pool. Utility was resource-specific: GSE306429 and MIX-Seq used cosine similarity between matched target and candidate response vectors; GDSC2 used negative absolute error on LN IC50. LINCS quantified the association between baseline similarity and matched-response concordance rather than task-level source-selection utility. Random-single, equal-weight ensemble, risk and abstention endpoints addressed separate decisions.

The analysis units differed by resource: LINCS used pair-condition records nested within cell pairs and matched perturbation conditions; GSE306429 used pair-condition records for association and target-perturbation tasks for utility; MIX-Seq used exact-time-matched tasks with perturbation-balanced resampling; and GDSC2 used target-drug tasks, drug-level means and crossed drug-by-target-model sensitivity analyses.

### Baseline expression and transcriptomic similarity

LINCS used GSE70138 Level 5 profiles. Baseline profiles were constructed from quality-controlled vehicle or DMSO and explicit untreated controls. Similarity between two cell lines was the Pearson correlation of their control-centroid profiles on the landmark-gene space after finite-value masking. Matched response profiles were compared by cosine similarity and supporting response metrics defined in the source analysis. Response-signature-strength covariates were included in the record-level association model.

GSE306429 used metadata-repaired limma response signatures at 24 hours over the MAIN gene space. Pair comparisons were restricted to exact compound-dose matches. The pairwise association estimand was the mean response cosine across shared conditions, while the task-level utility estimand compared the highest-similarity source with the mean eligible single-source utility. The two estimands were analyzed separately. For Fig. 3A, candidates were ranked by baseline similarity and graded relevance was defined as n minus each candidate’s average descending rank by observed task utility. NDCG@1 was the tie-aware DCG at rank 1, DCG@1 = q_(1)/log2(2), divided by the ideal DCG after sorting the relevance values; q_(1) is the tie-averaged relevance at the selected top position. This ranking diagnostic is distinct from the selected-minus-random ΔU endpoint.

MIX-Seq used the official cancer-derived background pool and required exact-time baseline availability for source eligibility. The primary endpoint was the most-similar single-source utility minus the random-single expectation. The weighted endpoint compared a similarity-weighted ensemble with an unweighted all-source ensemble. Missing baseline requests were logged and excluded without imputation, cross-time substitution or candidate recovery. The resource-level association, task-level utility and ensemble endpoints were kept separate because they use different targets, candidate pools and inferential units.

For the LINCS record-level model, response cosine similarity was regressed on baseline Pearson similarity and the Euclidean norms of the two matched response vectors, which were included as response-signature-strength covariates. The covariance was clustered by exact perturbagen-dose-time condition. Dyadic inference followed multiway-cluster methods (Cameron et al., 2011). The dyadic sensitivity analysis collapsed repeated records to 435 cell pairs and used two-way clustering on the two dyad endpoints together with node-label procedures. For GSE306429, the pair-level estimand was the mean response cosine across exact compound-dose matches, with shared-node-aware inference and leave-one-node-out diagnostics. In the task-level analyses, the highest-similarity candidate was compared with the mean utility across the eligible single-source pool.

### GDSC2 response and expression data

GDSC2 response data were taken from GDSC2_fitted_dose_response_27Oct23.xlsx and restricted to DATASET = GDSC2, NLME_RESULT_ID = 343 and WEBRELEASE = Y, with LN IC50 as the response endpoint (Yang et al., 2013). Baseline expression used the rnaseq_all_20241212.csv entry in rnaseq_all_20241212.zip with rsem TPM values. Exact SIDM model identity matching intersected 964 response models with 580 RNA-seq file models and yielded 396 analyzable joint models. The metadata-estimated overlap of 939 was retained as a resource-catalogue quantity and was not used as an analysis sample size.

After filtering all-missing and zero-variance genes, 37,233 genes remained. Baseline similarity was Pearson correlation across the processed expression matrix. The full analysis included 295 eligible drugs. Drug eligibility required at least 80 joint models, no duplicate records, at most one unique drug name and a valid screen signature. For each target model and drug, the target itself was excluded from the candidate pool and tasks with fewer than 20 eligible candidates were skipped. When multiple candidates shared the maximum Pearson similarity, the selected prediction was the mean LN IC50 across tied sources.

The response and expression tables were joined only through exact model identifiers; the catalogue-estimated overlap of 939 models was retained as a resource description and was not used as an analysis sample size. The primary full-resource estimate was the equal-weight mean of the 295 per-drug mean single-source utility differences (0.359). A task-weighted mean across the 97,077 observed target-drug tasks (0.363) was retained as a descriptive secondary statistic. The primary estimate and its percentile interval resampled drugs, not individual target-drug tasks.

### GDSC2 utility and aggregation endpoints

For GDSC2, candidate utility was U_t(c) = -abs(y_t - y_c), where y_t and y_c are the target and candidate LN IC50 responses, respectively.

Here y_t_ and y_c denote the target and candidate LN IC50 responses, respectively.

The selected-source endpoint was the utility of the highest-similarity source minus the mean utility across all eligible single-source candidates for the same target-drug task. The all-source combination endpoint compared the selected source with the equal-weight mean response across all eligible candidates. The primary ranking estimate was the equal-weight mean of 295 per-drug mean single-source utility differences (0.359); the task-weighted mean (0.363) was descriptive. For the selected-minus-ensemble endpoint, task-level paired differences were summarized within each drug and averaged equally across drugs; its point estimate was -0.083 and its 95% interval was obtained by drug bootstrap. The task-weighted selected-minus-ensemble mean (-0.081) was retained separately as descriptive and was not paired with that drug-bootstrap interval.

For target task t and eligible candidate set C_t, the ranking increment was Delta U_t = U_t(c*) minus the mean of U_t(c) over all c in C_t, where c* is the highest-similarity source. When several sources tied at maximum similarity, the selected prediction used their mean LN IC50. The aggregation endpoint compared the selected prediction with the equal-weight mean response across C_t and was evaluated separately from source ranking.

### Lineage-restricted GDSC2 analysis

The lineage-restricted analysis used the eligible subset of 92,324 target-drug tasks, 295 drugs, 377 target models and 18 represented tissues. The same-subset global comparator ranked all eligible candidate sources. The within-tissue comparator restricted candidate sources to the target tissue. The retention ratio was the within-tissue effect divided by the same-subset global effect. The policy endpoint compared global transcriptomic-similarity top-1 selection with tissue-matched random-source selection. Cancer-type sensitivity and candidate-level adjusted models were supporting analyses; they were not used to redefine the primary within-tissue estimand.

The lineage-restricted candidate pool was defined before evaluating lineage-conditioned outcomes. The same-subset global and within-tissue estimates therefore share the same target tasks and drug scope, while differing only in the source restriction. Coverage, cancer-type sensitivity and the adjusted candidate-level model were reported descriptively and did not alter the primary estimand.

### Statistical inference and resampling

LINCS record-level inference used condition-clustered covariance. Dyadic analyses accounted for shared cell-line endpoints with two-way clustering and node-label procedures. GSE306429 pair-level inference used shared-node-aware resampling and synchronous node-label permutations. MIX-Seq intervals used perturbation-balanced bootstrap resampling.

GDSC2 drug-bootstrap confidence intervals resampled the drug unit with replacement using 2,000 draws and seed 20260806 for the primary estimates. Crossed sensitivity intervals resampled drugs and target models with replacement using 2,000 draws and seed 20260807. Target-model-only sensitivity intervals held drugs fixed. Percentile intervals used the 2.5th and 97.5th percentiles. Identity-label sensitivity used 1,000 synchronous row and column model-identity permutations with seed 20260807 while holding response values, similarity values, eligibility and target self-exclusion fixed. Empirical permutation probabilities used the plus-one correction where implemented by the analysis code.

Permutation tests used the analysis-specific tail definition and plus-one empirical probabilities (Phipson & Smyth, 2010). Results at the attainable permutation floor are reported in the text as threshold values, with machine-precision values retained in the source tables. Perturbational intervals used target- or perturbation-balanced resampling as specified for each endpoint. The crossed sensitivity analysis resampled drugs and target models jointly; target-model-only intervals held the drug set fixed. All reported confidence intervals are percentile intervals.

### Use of AI-assisted tools

OpenAI Codex assisted with this final manuscript revision by checking statements against the supplied analysis code and source tables, checking consistency across the manuscript and Supporting Information, and editing the prose. It did not run the study analyses or generate or alter reported results in this revision. The authors remain responsible for the scientific interpretation and published content.

## Data and code availability

### Data availability

This study uses public LINCS GSE70138, GSE306429, MIX-Seq and GDSC2 resources; accessions, release identifiers and response filters are given in the Methods and Supporting Information. Derived tables supporting the figures and estimates are supplied as Supplementary Data and Source Data files. Code availability: A reviewer code package containing the analysis scripts, run-order notes and file hashes accompanies this submission. A public project-code repository URL or DOI is not available in this package.

## Declarations

### Ethics approval

not applicable to analyses of public de-identified datasets.

### Consent for publication

not applicable.

### Author contributions

H.Z. conceived and designed the study, developed the methodology, provided scientific supervision, interpreted the results, revised the manuscript and acquired funding. L.J. and B.L. performed analyses, curated data and maintained the analysis code. J.P. contributed to the original draft and interpretation. X.L. and X.W. curated data and prepared visualizations. All authors reviewed and approved the final manuscript.

## Acknowledgements

not applicable.

## References

Aguirre, M., Spence, J. P., Sella, G. & Pritchard, J. K. Gene regulatory network structure informs the distribution of perturbation effects. PLoS Computational Biology 21, e1013387 (2025). 10.1371/journal.pcbi.1013387

Ahlmann-Eltze, C., Huber, W. & Anders, S. Deep-learning-based gene perturbation effect prediction does not yet outperform simple linear baselines. Nature Methods 22, 1657–1661 (2025). 10.1038/s41592-025-02772-6

Barretina, J. et al. The Cancer Cell Line Encyclopedia enables predictive modelling of anticancer drug sensitivity. Nature 483, 603–607 (2012). 10.1038/nature11003

Ben-David, U. et al. Genetic and transcriptional evolution alters cancer cell line drug response. Nature 560, 325–330 (2018). 10.1038/s41586-018-0409-3

Bernett, J. et al. Critical evaluation of drug response prediction models with DrEval. Nature Communications 17, 4238 (2026). 10.1038/s41467-026-72903-w

Carli, F. et al. Learning and actioning general principles of cancer cell drug sensitivity. Nature Communications 16, 1654 (2025). 10.1038/s41467-025-56827-5

Cameron, A. C., Gelbach, J. B. & Miller, D. L. Robust inference with multiway clustering. Journal of Business & Economic Statistics 29, 238–249 (2011). 10.1198/jbes.2010.07136

Chawla, S. et al. Gene expression based inference of cancer drug sensitivity. Nature Communications 13, 7329 (2022). 10.1038/s41467-022-33291-z

Chen, J. et al. Deep transfer learning of cancer drug responses by integrating bulk and single-cell RNA-seq data. Nature Communications 13, 6494 (2022). 10.1038/s41467-022-34277-7

Costello, J. C. et al. A community effort to assess and improve drug sensitivity prediction algorithms. Nature Biotechnology 32, 1202–1212 (2014). 10.1038/nbt.2877

Crawford, J., Chikina, M. & Greene, C. S. Best holdout assessment is sufficient for cancer transcriptomic model selection. Patterns 5, 101115 (2024). 10.1016/j.patter.2024.101115

Csendes, G., Sanz, G., Szalay, K. Z. & Szalai, B. Benchmarking foundation cell models for post-perturbation RNA-seq prediction. BMC Genomics 26, 393 (2025). 10.1186/s12864-025-11600-2

DeMeo, B. et al. Active learning framework leveraging transcriptomics identifies modulators of disease phenotypes. Science 390, eadi8577 (2025). 10.1126/science.adi8577

Dixit, A. et al. Perturb-Seq: dissecting molecular circuits with scalable single-cell RNA profiling of pooled genetic screens. Cell 167, 1853–1866.e17 (2016). 10.1016/j.cell.2016.11.038

Feizi, N. et al. PharmacoDB 2.0: improving scalability and transparency of in vitro pharmacogenomics analysis. Nucleic Acids Research 50, D1348–D1357 (2022). 10.1093/nar/gkab1084

Garnett, M. J. et al. Systematic identification of genomic markers of drug sensitivity in cancer cells. Nature 483, 570–575 (2012). 10.1038/nature11005

Geeleher, P., Cox, N. & Huang, R. S. Clinical drug response can be predicted using baseline gene expression levels and in vitro drug sensitivity in cell lines. Genome Biology 15, R47 (2014). 10.1186/gb-2014-15-3-r47

Haibe-Kains, B. et al. Inconsistency in large pharmacogenomic studies. Nature 504, 389–393 (2013). 10.1038/nature12831

Iorio, F. et al. A landscape of pharmacogenomic interactions in cancer. Cell 166, 740–754 (2016). 10.1016/j.cell.2016.06.017

Keenan, A. B. et al. The Library of Integrated Network-Based Cellular Signatures NIH Program: system-level cataloging of human cells response to perturbations. Cell Systems 6, 13–24 (2018). 10.1016/j.cels.2017.11.001

Kernfeld, E., Yang, Y., Weinstock, J. S., Battle, A. & Cahan, P. A comparison of computational methods for expression forecasting. Genome Biology 26, 388 (2025). 10.1186/s13059-025-03840-y

Lamb, J. et al. The Connectivity Map: using gene-expression signatures to connect small molecules, genes, and disease. Science 313, 1929–1935 (2006). 10.1126/science.1132939

Lotfollahi, M., Wolf, F. A. & Theis, F. J. scGen predicts single-cell perturbation responses. Nature Methods 16, 715–721 (2019). 10.1038/s41592-019-0494-8

Lotfollahi, M. et al. Predicting cellular responses to complex perturbations in high-throughput screens. Molecular Systems Biology 19, e11501 (2023). 10.15252/msb.202211517

McFarland, J. M. et al. Multiplexed single-cell transcriptional response profiling to define cancer vulnerabilities and therapeutic mechanism of action. Nature Communications 11, 4296 (2020). 10.1038/s41467-020-17440-w

Pham, T.-H. et al. Chemical-induced gene expression ranking and its application to pancreatic cancer drug repurposing. Patterns 3, 100441 (2022). 10.1016/j.patter.2022.100441

Phipson, B. & Smyth, G. K. Permutation P-values should never be zero: calculating exact P-values when permutations are randomly drawn. Statistical Applications in Genetics and Molecular Biology 9, Article 8 (2010). 10.2202/1544-6115.1585

Qi, X. et al. Predicting transcriptional responses to novel chemical perturbations using deep generative model for drug discovery. Nature Communications 15, 9256 (2024). 10.1038/s41467-024-53457-1

Replogle, J. M. et al. Mapping information-rich genotype-phenotype landscapes with genome-scale Perturb-seq. Cell 185, 2559–2575.e28 (2022). 10.1016/j.cell.2022.05.013

Safikhani, Z. et al. Revisiting inconsistency in large pharmacogenomic studies. F1000Research 5, 83 (2017). 10.12688/f1000research.9611.3

Shimada, K., Hayano, M., Pagano, N. C. & Stockwell, B. R. Cell-line selectivity improves the predictive power of pharmacogenomic analyses and helps identify NADPH as biomarker for ferroptosis sensitivity. Cell Chemical Biology 23, 225–235 (2016). 10.1016/j.chembiol.2015.11.016

Srivatsan, S. R. et al. Massively multiplex chemical transcriptomics at single-cell resolution. Science 367, 45–51 (2020). 10.1126/science.aax6234

Subramanian, A. et al. A Next Generation Connectivity Map: L1000 platform and the first 1,000,000 profiles. Cell 171, 1437–1452.e17 (2017). 10.1016/j.cell.2017.10.049

Viñas Torné, R. et al. Systema: a framework for evaluating genetic perturbation response prediction beyond systematic variation. Nature Biotechnology 44, 1050–1059 (2026). 10.1038/s41587-025-02777-8

Warren, A. et al. Global computational alignment of tumor and cell line transcriptional profiles. Nature Communications 12, 22 (2021). 10.1038/s41467-020-20294-x

Yang, W. et al. Genomics of Drug Sensitivity in Cancer (GDSC): a resource for therapeutic biomarker discovery in cancer cells. Nucleic Acids Research 41, D955–D961 (2013). 10.1093/nar/gks1111

Zhang, N. et al. Predicting anticancer drug responses using a dual-layer integrated cell line-drug network model. PLoS Computational Biology 11, e1004498 (2015). 10.1371/journal.pcbi.1004498

